# Exploring the effect of capsaicin on gene expression and chemotherapy sensitization in gastric cancer cells

**DOI:** 10.1101/2024.09.04.611214

**Authors:** Weijian Meng, Sophia Xie, Rui Zhang, Jie Shen

**Author notes:** Corresponding author: Jie Shen, Tongji Hospital, Tongji Medical College, Huazhong University of Science and Technology, 1095 Jiefang Ave. Wuhan, Hubei 430030, China. These authors contribute equally to this work (Weijian Meng, Sophia Xie).

## Abstract

**Purpose:** Capsaicin has previously been demonstrated to exhibit anti-tumor effect in various cancer type. However, the effect of capsaicin on gene expression and its potential mechanism on chemotherapy sensitization were still uncertain.

**Method:** Human AGS gastric cancer cell line was treated with different concentrations of capsaicin, 5-fluorouracil and oxaliplatin. Cell viability was assessed using cell viability assay. High throughput RNA sequencing was used to screen differentially expressed genes triggered by capsaicin in AGS cells. qPCR and Western blotting were used to detect the expression of mRNAs and proteins induced by capsaicin.

**Result:** Capsaicin could significantly inhibit cell viability at a dose-depend manner in AGS gastric cancer cell line. Through high-throughput RNA sequencing, genes regulating DNA repair, DNA replication and chromosome assemble pathways were analyzed to be down-regulated by capsaicin. qPCR and western blot assay demonstrated that capsaicin could inhibit expression of the key enzymes (FEN1, LIG1 and PARP1) which play critical roles in DNA damage response and chemotherapy resistance. In vitro assay demonstrated that capsaicin could significantly induce chemo-sensitivity of 5-FU and Oxaliplatin at low dose.

**Conclusion:** Capsaicin could inhibit DNA repair pathway, which might contribute to cell growth inhibition and improvement of chemotherapy sensitization. These results revealed a novel function of capsaicin in DNA damage repair and provided new potential targets in cancer therapy.

## Introduction

Gastric cancer ranks third in cancer related death. Till now, 5-fluorouracil plus platinum agent is still the first line regimen in treatment of gastric cancer^1^. However, drug resistance is widely occurred thus limits its therapeutic efficiency^2^. Chemo-resistance could be caused by various mechanisms including: existence of cancer stem cells, hyperactivation of DNA repair pathways, dysregulation of apoptosis signals. Therefore, finding out an efficiency approach to induce chemo-sensitization is essential in cancer treatment.

Some natural dietary agents, which come from fruits, vegetables, or spices, exhibit potential therapeutic value in cancer treatment. Capsaicin, alternatively called trans-8-methyl-N-vanillyl-6-nonenamide, is the piquant component in red hot chili peppers which exhibits effects in cancer progression^3^. Previous researches have been demonstrated that capsaicin could inhibit cell proliferation in various cancer^4^.

Meanwhile, capsaicin was also reported to induce of apoptosis and cell cycle arrest. Some studies indicated that capsaicin could be used for cancer prevention^5^. However, the effect of capsaicin on gene expression and its potential mechanism on chemotherapy sensitization were still uncertain.

In this study, we found capsaicin could inhibit cell viability in AGS gastric cancer cell line. Through high-throughput RNA sequencing, genes regulating DNA repair, DNA replication and chromosome assemble pathways were analyzed to be down-regulated by capsaicin. qPCR assay and western blot demonstrated that capsaicin could inhibit expression of the key enzymes (FEN1, LIG1 and PARP1) which play critical roles in DNA damage response and chemotherapy resistance. In vitro assay showed that capsaicin could significantly induce chemo-sensitivity of 5-FU and Oxaliplatin at low dose. Our findings revealed a novel function of capsaicin in DNA damage repair and might provide new potential targets in cancer therapy.

## Methods

### Cell culture

Human gastric cancer cell line AGS was cultured in DMEM high-glucose culture medium containing 10%FBS, placed in a 5% CO2 incubator at 37 ^°^C, digested with 0.25% trypsin every 3 to 4 days, and passed through 1:3 cells with logarithmic growth phase for experiment.

### Cell viability assay

AGS cells were digested with trypsin, cell counts were carried out, and 1×10^4^/100μl cell suspension was prepared. 100μl per well was added into 96-well plates and cultured in cell incubators for 24h. After the cells were glued to the wall, drug treatment was administered. After two hours of incubation, the absorbance at 450 nm was measured using an enzyme label. Using the absorbance as a guide, the impact of medication treatment on cell activity was estimated.

### Western blot assay

Western blot was performed as previously described^6^. Total protein was extracted using NP40 lysate buffer supplemented with phenylmethylsulfonyl fluoride and cocktail protease inhibitor (MedChemExpress, Cat. #HY-K0010). The quantified protein samples were added to a 10% polyacrylamide gel (SDS-PAGE) and the proteins were separated by electrophoresis according to their molecular weight. The isolated proteins on the gel were transferred to a polyvinylidene fluoride membrane. The membrane is treated with a sealer containing a non-specific protein (skim milk powder) to close the non-specific binding sites on the membrane and reduce the background signal. The membrane is soaked in a solution containing a primary antibody specific to the target protein and incubated overnight, usually at 4°C. The membrane is soaked in 1:500 horseradish peroxidase-linked secondary antibodies (ABclonal, Cat. #AS014) and incubated for 2.5 h. Finally, the membranes were washed three times and visualized using an SuperKine™ West Femto Maximum Sensitivity Substrate kit (Abbkine, Cat. #BMU102). Primary antibodies used in western blot assay were as follows: PARP1 (ABclonal, Cat. #A0010); LIG1 (ABclonal, Cat. #A1858); FEN1 (ABclonal, Cat. #A1175); β -actin Rabbit mAb (ABclonal,Cat. #AC038).

### qPCR assay

The SteadyPure Universal RNA Extraction kit (ACCURATE, Cat. #AG21017) was used for total RNA extraction and reverse transcription was performed with HiScript II Q RT SuperMix for qPCR kit (Vazyme, Cat. #RR223-01) according to the manufacturer’s instructions. qPCR assay was performed using the ChamQ Universal SYBR qPCR Master Mix Kit (Vazyme, Cat.# Q711-02) in an ABI 7300Real-Time PCR System (Applied Biosystems, Foster City, CA, USA).GAPDH expression was used as an endogenous control, and the qPCR results were analyzed using the comparative Ct method (2^-ΔΔCt^). Primers used in qPCR assay were as follows:

LIG1:

Forward (5’-3’): GAAGGAGGCATCCAATAGCAG;

Reverse (5’-3’): ACTCTCGGACACCACTCCATT;

FEN1:

Forward (5’-3’): ATGACATCAAGAGCTACTTTGGC;

Reverse (5’-3’): GGCGAACAGCAATCAGGAACT;

PARP1:

Forward (5’-3’): CGGAGTCTTCGGATAAGCTCT;

Reverse (5’-3’): TTTCCATCAAACATGGGCGAC;

GAPDH:

Forward (5’-3’): TGTGGGCATCAATGGATTTGG;

Reverse (5’-3’): ACACCATGTATTCCGGGTCAAT.

### High throughput RNA sequencing

Total RNA was collected from AGS cells treated with capsaicin (at a dose of 250uM for 24h) or DMSO using TRIzol reagent according to the manufacturer’s protocol (n=3 per group). RNA was quantified using a NanoDrop ND-2000 (Thermo Scientific, USA), and RNA integrity was assessed using an Agilent Bioanalyzer 2100 (Agilent Technologies, USA). High throughput sequencing was performed by TsingKe biotech Co., Ltd. according to the manufacturers’ standard protocols.

The quality control and preliminary analysis of sequencing raw data was performed by Novogene biotech Co., Ltd. according to the standard pipeline. Gene expression level was measured by Fragments Per Kilobase of exon model per Million mapped fragments (FPKM). Differentially expressed genes were identified based on the fold change. The threshold set for upregulated and downregulated genes was a fold change more than 2 with p value < 0.01. Gene Ontology (GO) enrichment of the differentially expressed genes were performed using DAVID software (david.ncifcrf.gov). Protein-protein interaction analysis were performed using String program (string-db.org). The gene expression profile was analyzed using GSEA software (http://software.broadinstitute.org/gsea/).

### Statistical analysis

All experiments were repeated three times. (version 9.5.0 for Windows; GraphPad Software Inc., USA) was used for statistical analysis. All continuous data are presented as mean ± SD. Analysis of Variance (ANOVA) was used for statistical analysis and p<0.05 was set as statistical significance.

## Results

### Capsaicin inhibits cell viability in AGS gastric cancer cell line

Cell viability assay was used for detection the effect of capsaicin on cell viability in AGS gastric cancer cell line. AGS cells were treated with capsaicin at different concentrations (0.1μM, 1μM, 10μM, 25μM, 50μM, 100μM, 200μM, 300μM) for 24h, 48h and 72h respectively. The results showed that cell viability of AGS cells could be inhibited by capsaisin in a concentration-dependent manner (Fig. 1A-C). After 24h treatment, the IC50 of capsaicin for cell viability was calculated as 253.0μM (Fig. 1D).

**Fig 1.**
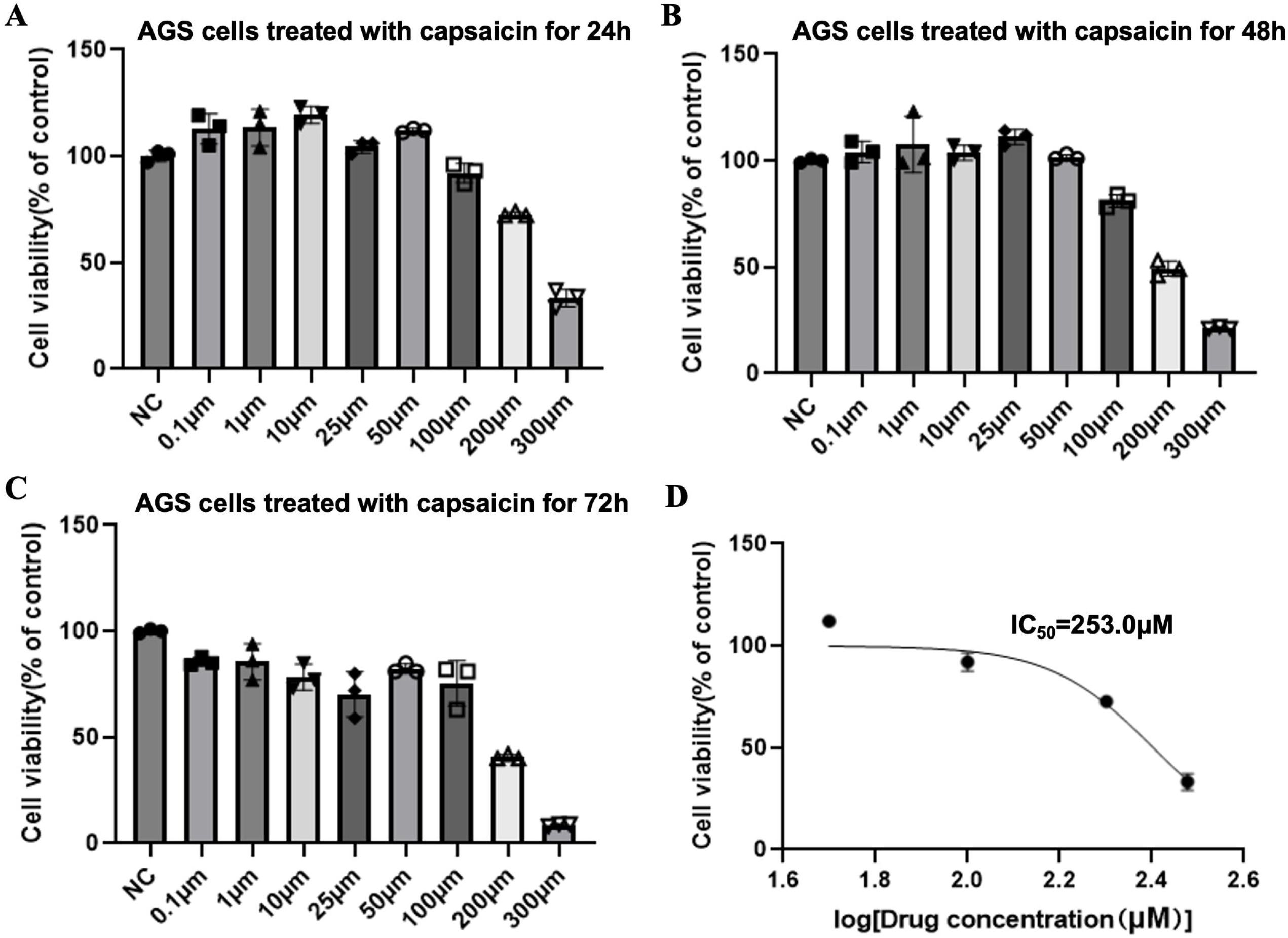
Capsaicin could inhibit cell viability in AGS cells. A. Cell viability of AGS cells after treating with capsaicin for 24h at different concentrations. DMSO was used as negative control. Histograms showed the mean ± SD of cell viability rate. B. Cell viability of AGS cells after treating with capsaicin for 48h at different concentrations. DMSO was used as negative control. Histograms showed the mean ± SD of cell viability rate. C. Cell viability of AGS cells after treating with capsaicin for 72h at different concentrations. DMSO was used as negative control. Histograms showed the mean ± SD of cell viability rate. D. AGS cells were treated with capsaicin with different concentrations for 24h. Then cell viability was measured and IC50 was calculated. IC50: half maximal inhibitory concentration.

### Capsaicin could inhibit DNA repair and chromosome assemble pathways in AGS cells

To elucidate the potential mechanisms of capsaicin mediated cell viability inhibition in gastric cancer cells, we collected mRNAs from AGS cells treated with either DMSO or capsaicin, and profiled gene expression using high-throughput sequencing. The differentially expressed gene (DEG) was defined as gene with expression fold change >2 and p<0.05. Base on this criterion, 1082 down-regulated DEGs and 1348 up-regulated DEGs in AGS cells treated with capsaicin were selected (Fig. 2A). Gene ontology (GO) enrichment was performed to explore the major function of these DEGs. We found that “chromosome segregation” and “DNA replication” occupied almost top ten GO categories of the downregulated DEGs in AGS cell treated with capsaicin (Fig. 2B). These results implied that capsaicin might down-regulate DNA replication and chromosome assemble genes in AGS cells.

**Fig 2.**
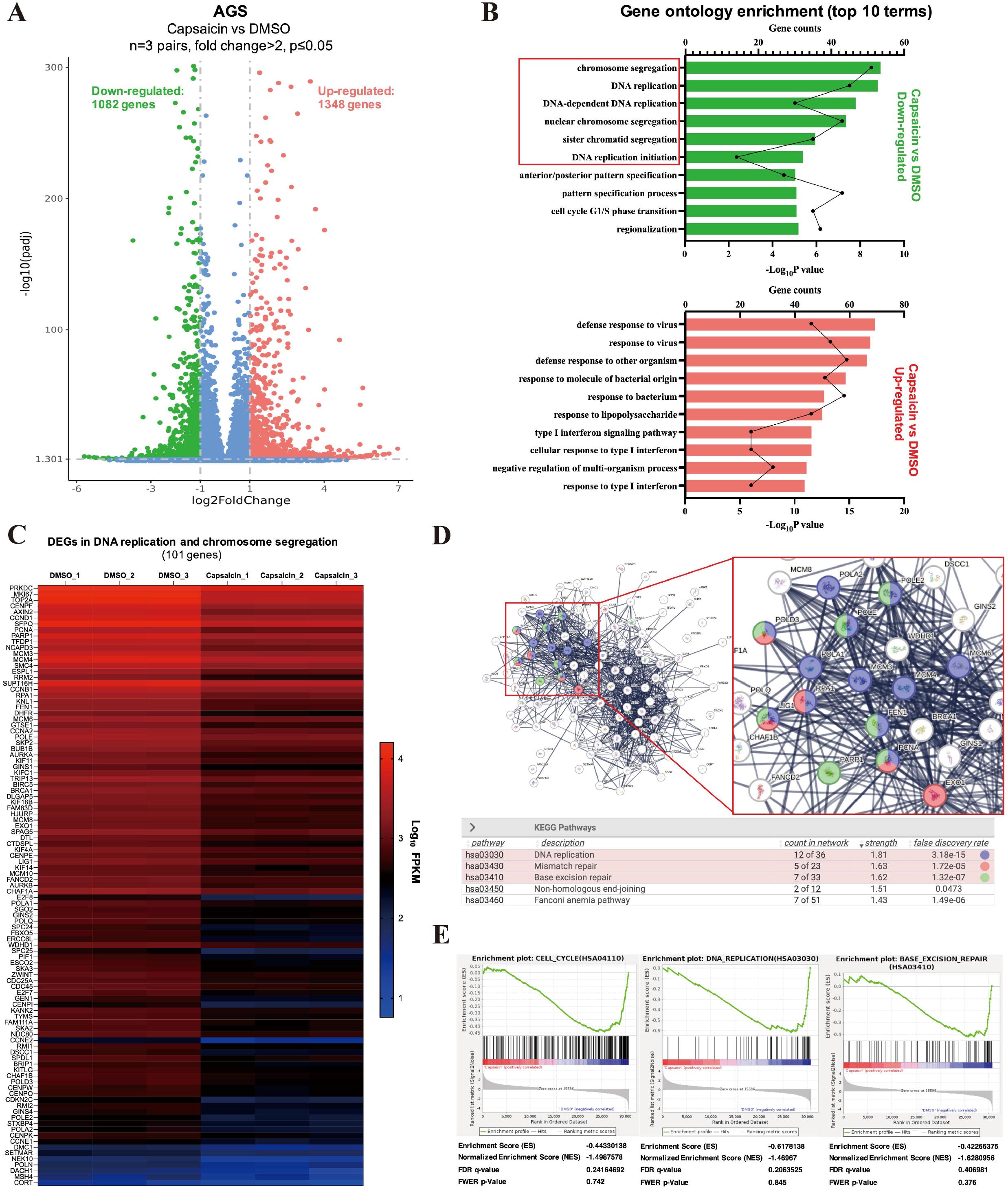
Capsaicin could inhibit DNA repair and chromosome assemble pathways in AGS cells. A. Volcano plot diagram of gene expression in AGS cell with control or capsaicin treatment. AGS cells were treated with capsaicin at a dose of 250uM for 24h (DMSO was used as control), then high throughput sequencing was performed (N=3 pairs). The differentially expressed gene (DEG) was defined as gene with expression fold change >2 and p<0.05. B. Top-10 Gene ontology (GO) enrichment categories of the up/down-regulated DEGs in AGS cells. Green, GO enrichment categories of down-regulated DEGs in capsaicin vs. DMSO; red, GO enrichment categories of up-regulated DEGs in capsaicin vs. DMSO; bar: Log10 p value; dot, gene counts. C. Heatmap presented 101 DEGs, which were classified in “chromosome segregation”, “DNA replication”, “DNA-dependent DNA replication”, “nuclear chromosome segregation”, “sister chromatid segregation” and “DNA replication initiation” categories. FPKM, fragments per kilobase of exon model per million mapped fragments. D. String program was used to annotate the functions and potential interactions of these 101 DEGs. The potential downstream of capsaicin in MMR (POLD3, RPA1, LIG1, PCNA, EXO1) and BER (POLE2, POLE, POLD3, LIG1, FEN1, PARP1, PCNA) were also identified. E. Gene set enrichment analysis (GSEA) of AGS cell with control (DMSO) or capsaicin treatment (n = 3 pairs). KEGG subset of “C2: curated gene sets” was used in this analysis. The normalized p values of the presented gene sets were less than 0.01. ES: enrichment score; NES: normalized enrichment score; FDR-q: false discovery rate q value; FWER-p: family-wise error rate p value.

Next, we focused on the DEGs in the DNA replication and chromosome segregation categories, and identified 101 down-regulated genes in AGS cells treated with capsaicin (Fig. 2C). Using String program (string-db.org), we found that the function of these genes could be significantly enriched in mismatch repair (MMR) (rank 2) and base excision repair (BER) (rank 3) pathways (Fig. 2D). String analysis also identified the potential downstream genes of capsaicin in MMR (POLD3, RPA1, LIG1, PCNA, EXO1) and BER (POLE2, POLE, POLD3, LIG1, FEN1, PARP1, PCNA) pathways (Fig. 2D). To further verify whether capsaicin could regulate cell cycle and DNA repair in AGS cells, gene set enrichment analysis was performed. Our results revealed that, compared with capsaicin treatment group, cell cycle; DNA replication; and base excision repair could be enriched in control group with statistical significance (normalized p value < 0.01) (Fig. 2E). These evidence indicated that capsaicin could inhibit DNA repair in BER dependent manner and negatively regulated cell proliferation in gastric cancer cells.

### Capsaicin suppresses LIG1, PARP1 and FEN1 expression in AGS cells

End-polishing by flap endonuclease 1 (FEN1), the DNA ligases 1 (LIG1) are the key enzymes in BER pathway, while poly ADP-ribose polymerase (PARP1) is essential in DNA repair and maintaining genomic integrity.^7,8^ Based on the high-hroughput sequencing data, capsaicin had potential regulatory effect of FEN1, LIG1 and PARP1, thus, these three genes were selected and validated. qPCR and Western blot were used to detect the expression of RNA and protein levels after treatment with different concentrations of capsaicin (50μM, 100μM, 253μM) for 24h, respectively. The results showed that capsaicin significantly inhibited the expression of LIG1, PARP1 and FEN1 at both RNA and protein levels in AGS cells (Fig. 3A, B).

**Fig 3.**
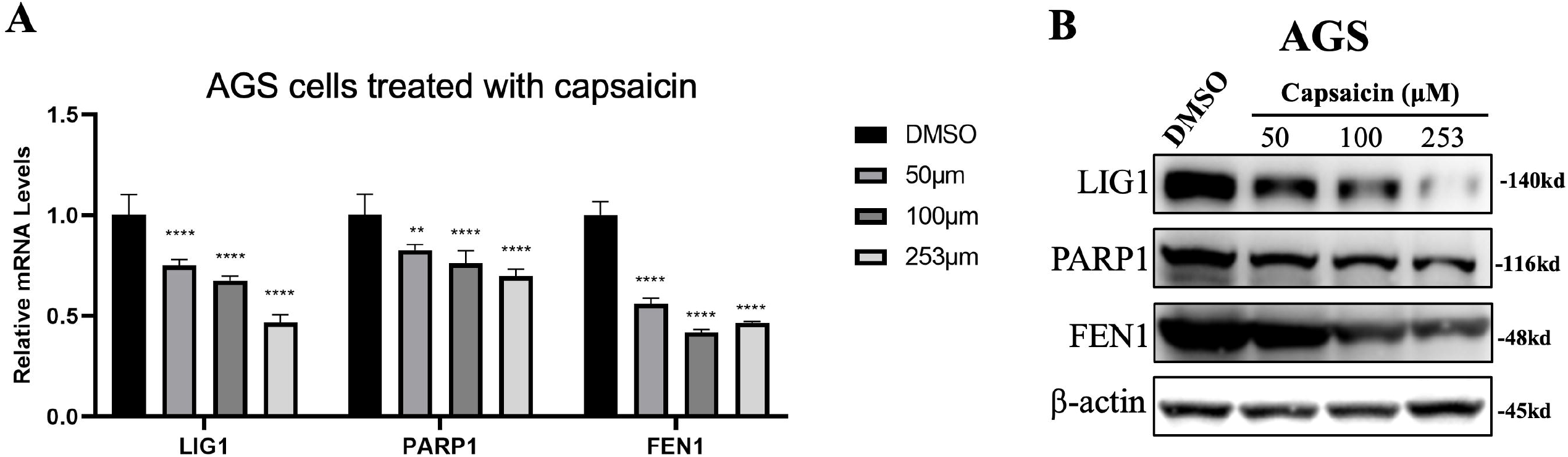
Capsaicin suppresses LIG1, PARP1 and FEN1 expression in AGS cells. A. AGS cells were treated with DMSO or capsaicin (50μM,100μM,253μM) for 24h, then qPCR was performed to detect the expression level of LIG1, PARP1 and FEN1. GAPDH was used as endogenous control. Three duplications were performed in each experiment. Data are represented as mean ± SD. **p < 0.01, ****p < 0.0001 by ANOVA. B. Cell lysates from AGS cells treated with DMSO or capsaicin(50μM,100μM,253μM) for 24h, then immunoblot was performed. Lysates were probed for LIG1, PARP1, FEN1 antibodies and β-actin was used as a loading control.

### Capsaicin could induce drug sensitivity of 5-FU and oxaliplatin in AGS cells

DNA damage response and DNA repair were essential in drug resistance of chemotherapy. To further validate whether capsaicin could affect chemo-sensitivity in gastric cancer cells, we first treated AGS cells with 5-Fluorouracil (5-FU) at different concentration and found the IC50 of 5-FU was 343.2μM (Fig. 4A). Interesting, combine treating with low dose of capsaicin (50μM), the 5-FU IC50 in AGS decreased to 195.2μM (Fig. 4B). Similarly, treating with low dose of capsaicin, the IC50 of Oxalipatin in AGS cells decreased from 54.2μM to 27.6μM (Fig. 4C, D). The results indicated that capsaicin could enhance the chemo-sensitivity of 5-FU and Oxalipatin in gastric cancer cells.

**Fig 4.**
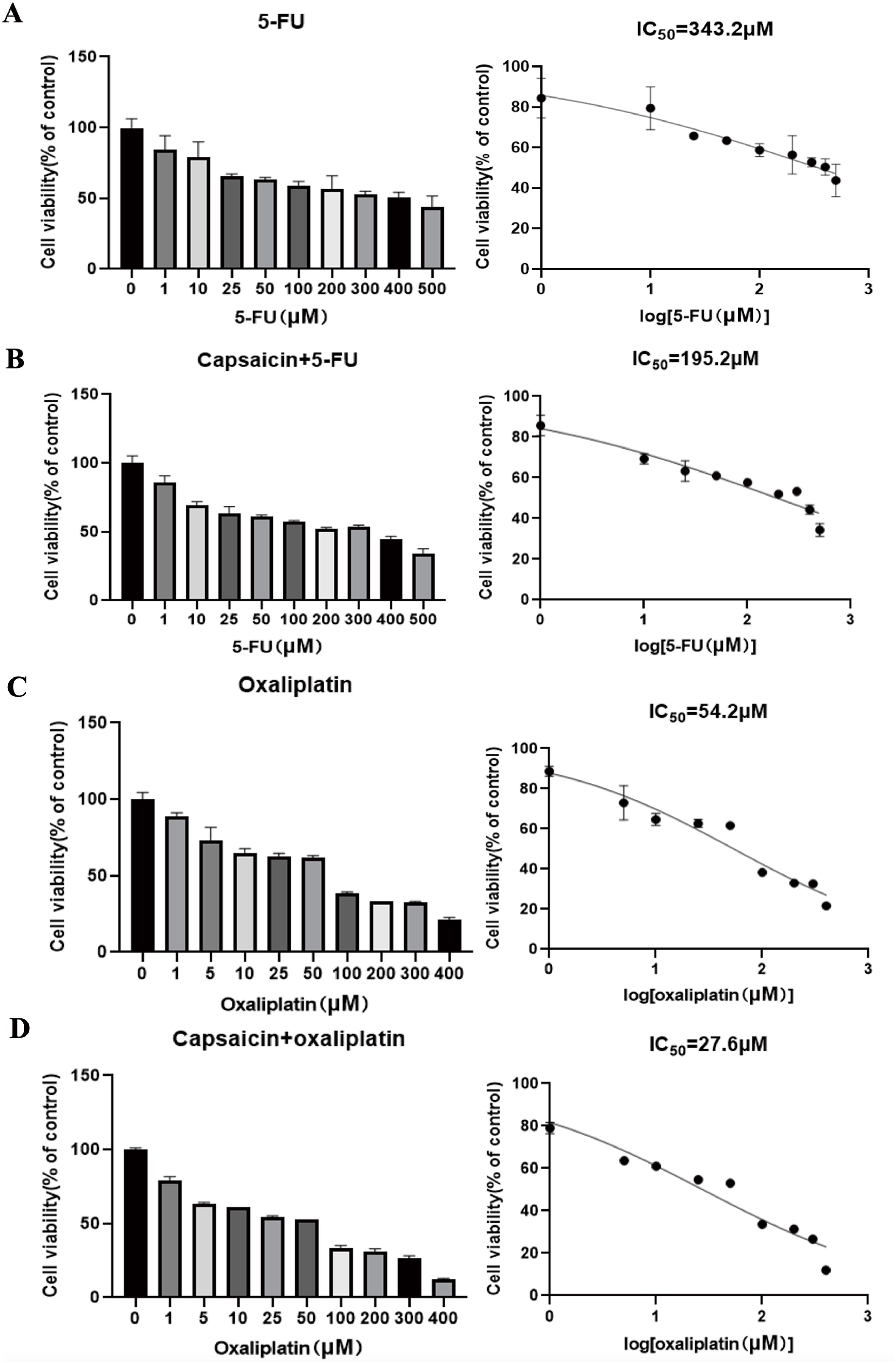
Capsaicin could induce drug sensitivity of 5-FU and oxaliplatin in AGS cells. A. AGS cells were treated with different concentrations of 5-FU for 24 hours, then cell viability assay was performed to detect cell viability and calculate IC50 of 5-FU. DMSO was used as negative control. B. AGS cells were co-treated with capsaicin at a dose of 50μM and different concentrations of 5-FU for 24 hours. Then cell viability assay was performed to detect cell viability and calculate IC50 of 5-FU. DMSO was used as negative control. C. AGS cells were treated with different concentrations of Oxaliplatin for 24 hours, then Cell viability assay was performed to detect cell viability and calculate IC50 of Oxaliplatin. DMSO was used as negative control. D. AGS cells were co-treated with capsaicin at a dose of 50μM and different concentrations of Oxaliplatin for 24 hours. Then Cell viability assay was performed to detect cell viability and calculate IC50 of Oxaliplatin. DMSO was used as negative control.

## Discussion

Previous studies unveiled an epidemiological evidence that spicy diet exhibited effects against cancer initiation and progression^9^. Among the phytochemical components of the spicy food, capsaicin has been demonstrated to play an important role in regulation of cell survival, growth arrest, angiogenesis and tumor metastasis^3^. Several researches evaluating the effect of capsaicin, and found capsaicin could increase of cell-cycle arrest and induce apoptosis in various cancer types^10^. Although the potential anti-tumor characteristics of capsaicin exhibited a good application prospect in cancer therapy, the effect of capsaicin on gene expression and its mechanism on chemotherapy sensitization were still uncertain.

In this current research, we provided evidence that capsaicin could significantly inhibit cell viability and proliferation in gastric cancer cells. Through high throughput sequencing, we found capsaicin could significantly inhibited the expression of genes, which were involved in DNA replication, especially in base excision repair pathway. Base excision repair is the main pathway for the repair of small base lesions in the DNA damage response of mammalian cells^11^. Briefly, BER is initiated by DNA glycosylases which recognize and excise the aberrant base, then AP endonuclease 1 (APE1) recognizes these abasic sites, incises DNA backbone and cooperates with DNA polymerases to inserts one or more nucleotides^8^. Following end-polishing by FEN1, the DNA ligases (e.g., LIG1, LIG3) reconstitute the integrity of the DNA backbone and complete BER process. Our data demonstrated that capsaicin could downregulated the expression level of FEN1 and LIG1 in AGS cells. This may provide a new insight on the findings which capsaicin could block BER pathway and induce cell cycle arrest in tumor cells. The mechanism of how capsaicin regulated FEN1/LIG1 expression still need to be elucidated in further studies.

5-Fluorouracil plus platinum agent (oxalipatin) is still the first-line chemotherapy for gastric cancer^1^. 5-FU induces cytotoxicity by inhibiting thymidylate synthase and disrupting essential biosynthetic processes in DNA replication, while platinum forms intra-/inter-strand crosslinks with DNA thus leading to the inhibition of DNA synthesis^12,13^. With a widespread usage of these agents, an increasing number of drug resistance have emerged^2^. Therefore, overcoming chemotherapy resistance has become crucial in cancer therapy. Evidences showed that DNA lesions associated with 5-fluorouracil therapy could be repaired by BER^14^, meanwhile FEN1 could block platinum chemo-sensitization and synthetic lethality through suppressing double-strand breaks accumulation^15^. Our study unveiled a novel function of capsaicin in inhibiting FEN1 and BER expression pathway. In vitro assay also demonstrated that capsaicin could significant induce cyto-toxicity of 5-FU and Oxaliplatin in gastric cancer cell line. These results indicated a potential pharmacobiology application of capsaicin in anti-cancer chemotherapy.

Another interesting finding in our study is that capsaicin could act as a potential inhibitor of PARP1 expression. PARP1 is a well-studied enzyme in the response to DNA damage by loosening chromatin at specific sites to promote DNA repair and maintain genomic integrity^7^. Due to the central role in DNA damage repair cascade, PARP1 inhibition is significantly more toxic in cancer cells with homologous recombination (HR) deficiency than in normal cells^16^. Currently, PARP inhibitors (PARPi), such as Olaparib, Rucaparib Niraparib and Talazoparib, have been emerged as a therapeutic strategy for tumors with high genomic instability^17^. However, artificial PARPi frequently exhibits hematological and gastrointestinal side-effects, meanwhile, multiple mechanisms (e.g., restoration of HR repair, alternative DNA repair pathways, decreased PARPi pharmacodynamic effect) contributing to PARPi resistance limit its usage in clinical practice^18^. In this study, we found that Capsaicin presents a latent capacity to inhibit both PARP1 expression and BER pathway. Thus, capsaicin may act as a new breakthrough in anti-tumor therapeutic strategy.

Through RNA sequencing data, we also found that up-regulated genes triggered by capsaicin could enrich in immune response signals, such as cell defense to virus, cell response to bacterium and type-I interferon pathways. Likewise, the receptor of capsaicin, TRPV1, has been reported to induce pro-inflammatory cytokine production when activated^19^. Till now, researches on the immunological regulation of TRPV1 is focused on neuro-immune interactions^20^. The effect of capsaicin/TRPV1 in tumor immunology is still uncertain. Our data provided a new hypothesis that capsaicin could play an effect on tumor immune response in gastric cancer. Further research on the characteristics and function of capsaicin in regulating tumor immunology is required.

In conclusion, our study demonstrated that capsaicin could inhibit DNA repair, thereby inhibited cell viability and improved the sensitivity of chemotherapy in gastric cancer cells. These findings revealed the biological function of capsaicin in tumor suppression and provided potential targets in cancer therapy.

## Funding

This work was supported by Hubei Provincial Natural Science Foundation of China (Grant numbers: 2023 AFB118)

## Acknowledgements

We would like to thank the Experimental Medicine Center of Tongji Hospital for providing support to our experiments.

## Author Contributions

J.S. was responsible for study design and project administration. W.M. and S.X. were responsible for performing experiments and data collecting; W.M, S.X. and R.Z. contributed to extracting and analyzing data; W.M. and S.X. wrote the manuscript; J.S. revised and edited manuscript; J.S. contributed to fu nding acquisition.

## Competing Interests

The authors declare that they have no conflict of interest.

## Data availability

All data generated or analyzed during this study are included in this article. The datasets used and/or analyzed during the current study are available from the first author and corresponding author upon reasonable request.

## Notes

### Competing Interest Statement

The authors have declared no competing interest.

## Reference

1. Yamashita, K., Hosoda, K., Niihara, M., and Hiki, N. (2021). History and emerging trends in chemotherapy for gastric cancer. Ann Gastroenterol Surg 5, 446–456. 10.1002/ags3.12439.

2. Liu, J., Yuan, Q., Guo, H., Guan, H., Hong, Z., and Shang, D. (2024). Deciphering drug resistance in gastric cancer: Potential mechanisms and future perspectives. Biomed Pharmacother 173, 116310. 10.1016/j.biopha.2024.116310.

3. Chapa-Oliver, A.M., and Mejia-Teniente, L. (2016). Capsaicin: From Plants to a Cancer-Suppressing Agent. Molecules 21. 10.3390/molecules21080931.

4. Hacioglu, C. (2022). Capsaicin inhibits cell proliferation by enhancing oxidative stress and apoptosis through SIRT1/NOX4 signaling pathways in HepG2 and HL-7702 cells. J Biochem Mol Toxicol 36, e22974. 10.1002/jbt.22974.

5. Radhakrishna, G.K., Ammunje, D.N., Kunjiappan, S., Ravi, K., Vellingiri, S., Ramesh, S.H., Almeida, S.D., Sireesha, G., Ramesh, S., Al-Qahtani, S., et al. (2024). A Comprehensive Review of Capsaicin and Its Role in Cancer Prevention and Treatment. Drug Res (Stuttg) 74, 195–207. 10.1055/a-2309-5581.

6. Feng, S., Huang, Q., Deng, J., Jia, W., Gong, J., Xie, D., Shen, J., and Liu, L. (2022). DAB2IP suppresses tumor malignancy by inhibiting GRP75-driven p53 ubiquitination in colon cancer. Cancer Lett 532, 215588. 10.1016/j.canlet.2022.215588.

7. Sinha, S., Molla, S., and Kundu, C.N. (2021). PARP1-modulated chromatin remodeling is a new target for cancer treatment. Med Oncol 38, 118. 10.1007/s12032-021-01570-2.

8. Visnes, T., Grube, M., Hanna, B.M.F., Benitez-Buelga, C., Cazares-Korner, A., and Helleday, T. (2018). Targeting BER enzymes in cancer therapy. DNA Repair (Amst) 71, 118–126. 10.1016/j.dnarep.2018.08.015.

9. Aggarwal, B.B., and Shishodia, S. (2006). Molecular targets of dietary agents for prevention and therapy of cancer. Biochem Pharmacol 71, 1397–1421. 10.1016/j.bcp.2006.02.009.

10. Szallasi, A. (2023). Capsaicin and cancer: Guilty as charged or innocent until proven guilty? Temperature (Austin) 10, 35–49. 10.1080/23328940.2021.2017735.

11. Curtin, N.J. (2012). DNA repair dysregulation from cancer driver to therapeutic target. Nat Rev Cancer 12, 801–817. 10.1038/nrc3399.

12. Sethy, C., and Kundu, C.N. (2021). 5-Fluorouracil (5-FU) resistance and the new strategy to enhance the sensitivity against cancer: Implication of DNA repair inhibition. Biomed Pharmacother 137, 111285. 10.1016/j.biopha.2021.111285.

13. Zhang, C., Xu, C., Gao, X., and Yao, Q. (2022). Platinum-based drugs for cancer therapy and anti-tumor strategies. Theranostics 12, 2115–2132. 10.7150/thno.69424.

14. Vodenkova, S., Jiraskova, K., Urbanova, M., Kroupa, M., Slyskova, J., Schneiderova, M., Levy, M., Buchler, T., Liska, V., Vodickova, L., et al. (2018). Base excision repair capacity as a determinant of prognosis and therapy response in colon cancer patients. DNA Repair (Amst) 72, 77–85. 10.1016/j.dnarep.2018.09.006.

15. Mesquita, K.A., Ali, R., Doherty, R., Toss, M.S., Miligy, I., Alblihy, A., Dorjsuren, D., Simeonov, A., Jadhav, A., Wilson, D.M., 3rd, et al. (2021). FEN1 Blockade for Platinum Chemo-Sensitization and Synthetic Lethality in Epithelial Ovarian Cancers. Cancers (Basel) 13. 10.3390/cancers13081866.

16. Farmer, H., McCabe, N., Lord, C.J., Tutt, A.N., Johnson, D.A., Richardson, T.B., Santarosa, M., Dillon, K.J., Hickson, I., Knights, C., et al. (2005). Targeting the DNA repair defect in BRCA mutant cells as a therapeutic strategy. Nature 434, 917–921. 10.1038/nature03445.

17. Bohi, F., and Hottiger, M.O. (2024). Expanding the Perspective on PARP1 and Its Inhibitors in Cancer Therapy: From DNA Damage Repair to Immunomodulation. Biomedicines 12. 10.3390/biomedicines12071617.

18. Mitri, Z., Goodyear, S.M., and Mills, G. (2024). Strategies for the prevention or reversal of PARP inhibitor resistance. Expert Rev Anticancer Ther, 1-17. 10.1080/14737140.2024.2393251.

19. Socala, K., Jakubiec, M., Abram, M., Mlost, J., Starowicz, K., Kaminski, R.M., Ciepiela, K., Andres-Mach, M., Zagaja, M., Metcalf, C.S., et al. (2024). TRPV1 channel in the pathophysiology of epilepsy and its potential as a molecular target for the development of new antiseizure drug candidates. Prog Neurobiol 240, 102634. 10.1016/j.pneurobio.2024.102634.

20. Maximiano, T.K.E., Carneiro, J.A., Fattori, V., and Verri, W.A. (2024). TRPV1: Receptor structure, activation, modulation and role in neuro-immune interactions and pain. Cell Calcium 119, 102870. 10.1016/j.ceca.2024.102870.

